# A Bayesian framework for inter-cellular information sharing improves dscRNA-seq quantification

**DOI:** 10.1101/2020.04.10.035899

**Authors:** Avi Srivastava, Laraib Malik, Hirak Sarkar, Rob Patro

## Abstract

**Motivation:** Droplet based single cell RNA-seq (dscRNA-seq) data is being generated at an unprecedented pace, and the accurate estimation of gene level abundances for each cell is a crucial first step in most dscRNA-seq analyses. When preprocessing the raw dscRNA-seq data to generate a count matrix, care must be taken to account for the potentially large number of multi-mapping locations per read. The sparsity of dscRNA-seq data, and the strong 3’ sampling bias, makes it difficult to disambiguate cases where there is no uniquely mapping read to any of the candidate target genes.

**Results:** We introduce a Bayesian framework for information sharing across cells within a sample, or across multiple modalities of data using the same sample, to improve gene quantification estimates for dscRNA-seq data. We use an anchor-based approach to connect cells with similar gene expression patterns, and learn informative, empirical priors which we provide to alevin’s gene multi-mapping resolution algorithm. This improves the quantification estimates for genes with no uniquely mapping reads (i.e. when there is no unique intra-cellular information). We show our new model improves the per cell gene level estimates and provides a principled framework for information sharing across multiple modalities. We test our method on a combination of simulated and real datasets under various setups.

**Availability:** The information sharing model is included in alevin and is implemented in C++14. It is available as open-source software, under GPL v3, at https://github.com/COMBINE-lab/salmon as of version 1.1.0.

## 1 Introduction

RNA sequencing, with subsequent gene and transcript quantification, has been an important tool for exploring genome-wide expression patterns using both bulk and single-cell experiments. With recent advancements in single-cell transcriptomic sequencing technologies, various droplet-based RNA sequencing (dscRNA-seq) methods (Macosko *et al.*, 2015; Klein *et al.*, 2015; Zheng *et al.*, 2017) have gained popularity due to their ability to generate data with higher quantitative accuracy, sensitivity, and throughput than previous approaches. These dscRNA-seq protocols have a unique output where each read is associated with a cell barcode, to facilitate separation of information between individual cells, and a unique molecular identifier (UMI) tag, that allows detecting and deduplicating PCR amplified molecules. Multiple preprocessing pipelines exist that use varying algorithms and methodologies to perform cell barcode correction and whitelisting, read alignment or mapping, and UMI deduplication, to eventually provide gene quantification estimates for each cell. Some of these pipelines use complete alignment of the reads to the reference, such as alevin (Srivastava *et al.*, 2019), STARsolo (Dobin, 2019), Cell-Ranger (Zheng *et al.*, 2017) and Hera-T (Tran *et al.*, 2019) whereas others use lightweight mapping methods, such as bustools (Melsted *et al.*, 2019). To the best of our knowledge, each method, except alevin, discards reads that multi-map between genes. To date, such approaches validate accuracy by demonstrating near-perfect correlation to estimates from Cell-Ranger.

In alevin, Srivastava *et al.* (2019) propose a novel framework for generating accurate gene-expression estimates for each cell given the read sequences from a dscRNA-seq experiment. It is shown how discarding gene multi-mapping reads, as is typically done by other existing dscRNA-seq quantification pipelines, can lead to biased and inaccurate expression estimates for certain genes and gene families. Subsequently, it is also demonstrated that alevin reduces this bias by providing a framework for assigning multi-mapping reads to genes rather than discarding them. Specifically, after a UMI resolution and deduplication phase (which assigns multi-mapping UMIs on the basis of parsimony), UMIs are placed into gene-level equivalence classes, associating each UMI with the set of genes to which it maps. Ambiguous reads that belong to equivalence classes with more than one gene label are probabilistically-assigned using an expectation-maximization (EM) algorithm. The EM algorithm works by integrating information from reads that are confidently assigned to a single gene, either as a result of the parsimony-based UMI resolution algorithm or because this was the only gene to which the underlying read aligns. This information helps to disambiguate reads that belong to multi-gene equivalence classes, and it is shown, through various analyses, that the framework provides better gene expression estimates than approaches that discard multi-mapping reads.

However, in situations where there is no unique evidence to disambiguate and assign a read among genes from its equivalence class with some confidence, the optimization method used by alevin uniformly divides the read count across all genes from the single equivalence class. This set of genes is then labeled as tier 3 in the alevin output. Genes within a cell that have some unique evidence, or share equivalence class with genes that do, are labeled as tier 2. Hence, tier 2 genes are assigned read counts with some level of confidence by the EM algorithm. Finally, tier 1 contains genes that have reads uniquely assigned to them at the UMI deduplication step, and hence their count can be estimated with the greatest confidence by the EM algorithm. This method of equivalence class and tier assignment is further detailed in Figure 1. In this study, we focus on genes labeled as tier 3, and propose an approach for improving the accuracy of their quantification, instead of uniformly dividing read counts between them.

**Fig. 1:**
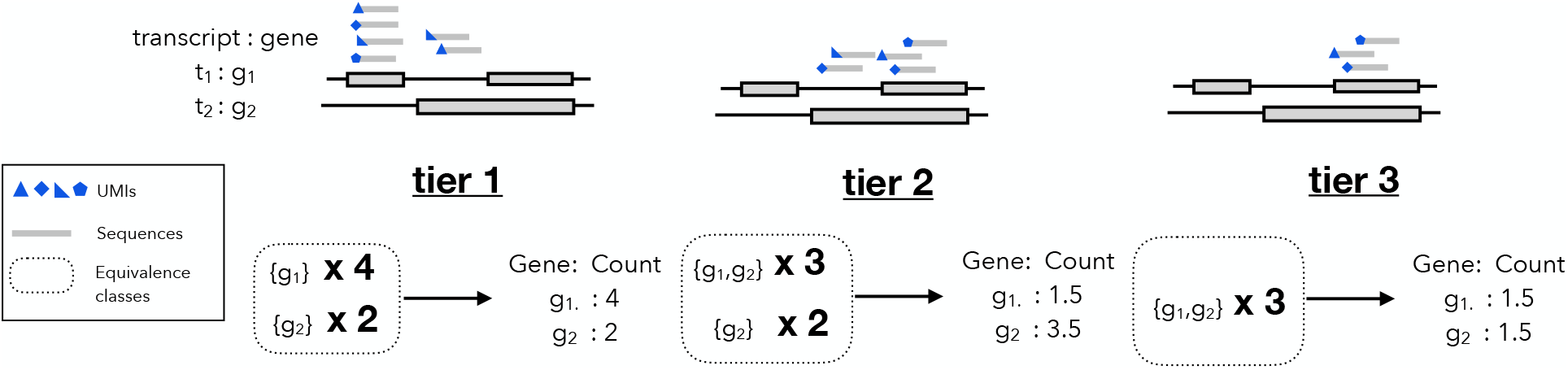
Alevin categorizes the quantification estimates based on their confidence into three tiers. Here we assume two transcript *t*_1_ and *t*_2_ coming from two genes *g*_1_ and *g*_2_ respectively. The toy example shows three tiers (from left to right) based on the mapping of the reads. Tier 1 estimates are from gene unique equivalence classes, tier 2 estimates are the cases where any gene in the equivalence class has unique evidence and finally tier 3 when no gene in the equivalence class has uniquely mapping reads.

Our proposed model works by sharing information, either across closely-related cells within the sample, or derived in some other fashion from the assay, such as in the case of spatial transcriptomics data. This information is integrated into the inference algorithm by introducing empirical Bayesian priors, and we show that the proposed Bayesian framework improves gene abundance estimates for tier 3 genes under various metrics, based on tests using simulated and real datasets in different setups. The idea of sharing information across data modalities, using an empirical prior, has been previously considered in the context of bulk RNA-seq (Liu *et al.*, 2016). Relatedly, the idea of sharing information across samples has also been applied in the context of imputation for various types of sparse genomic datasets, such as SNP genotyping and GWAS studies (Chou *et al.*, 2016; Visscher *et al.*, 2017). However, for single-cell quantification data, most imputation methods rely on intrinsic properties of the data due the absence of an external reference and work only *post hoc* on already generated gene count matrices (Tang *et al.*, 2018; Huang *et al.*, 2018; Wang *et al.*, 2019; Li and Li, 2018; Miao *et al.*, 2019; Chen and Zhou, 2018; Gong *et al.*, 2018; Van Dijk *et al.*, 2018; Wagner *et al.*, 2017; Talwar *et al.*, 2018; Eraslan *et al.*, 2019; Arisdakessian *et al.*, 2019; Amodio *et al.*, 2019; Deng *et al.*, 2019; Lopez *et al.*, 2018; Linderman *et al.*, 2018; Mongia *et al.*, 2019; Zhang and Zhang, 2018). Therefore, they do not have access to either the information contained in, or the constraints imposed by, the UMI-to-gene mappings. Our approach, on the other hand, utilizes shared information directly in the quantification phase to improve UMI assignment and resolution of multi-mapping reads. Furthermore, this information is used only in the form of an empirical prior, and the resulting quantification estimates are still strictly constrained by the observed data. Hence, the likelihood of inducing globally significant false signals, as has been reported in the case of some single-cell RNA-seq imputation methods (Andrews and Hemberg, 2018), is small.

## 2 Methods

### 2.1 Bayesian framework

After UMI deduplication, alevin models the read assignment problem as an optimization problem and iteratively assigns the ambiguous reads to potential candidates in a manner that maximizes (at least locally, within a cell) the joint likelihood. However, it cannot utilize the confidence information from neighboring cells, or from cells of the same type. Since a high level of sparsity is an inherent property of contemporary dscRNA-seq experiments (Hicks *et al.*, 2018), and due to the random process of capturing RNA molecules, in expectation, sampling can exhibit considerable variation across cells. Hence, we expect cells in an experiment to fall into categories of specific cell-types, and for cells of the same type to share similar expression patterns (Stuart *et al.*, 2019). However, for a specific gene, we do not expect that the molecules originating from the gene will be uniformly captured and sampled equally well across all cells of the type. Therefore, sharing confidence in the expression estimates across cells can be particularly effective in improving cell-level expression estimation. Similarly, we expect information from other assays, using either the same cells or even the same cell-type, to exhibit highly correlated gene abundances. We integrate this information using Bayesian priors by changing our optimization algorithm from an expectation maximization (EM) algorithm to a Variational Bayesian optimization algorithm (VBEM) (Nariai *et al.*, 2013) with an informative prior for low-information genes, i.e. genes assigned to tier 3. This is a variant of the same collapsed Variational Bayesian estimation method used in Salmon (Patro *et al.*, 2017) for bulk RNA-seq abundance estimation.

Similar to Salmon’s VBEM, we aim to quantify the expression, given a set of known genes 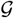 and a set of gene-level equivalence classes *ε* with their associated UMIs. Each equivalence class is labeled with a set of genes and has an associated set of UMIs, such that each UMI is attributed to at least one read that multi-maps only across the set of genes in the equivalence class. Here the set of UMIs are taken after appropriate deduplication using alevin’s graph-based UMI deduplication algorithm (Srivastava *et al.*, 2019). We use the VBEM algorithm to allow sharing quantification information across cells via the use of priors. Specifically, we define the gene-UMI count assignment matrix as 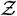, where, based on *ε*, *z*_*ij*_ = 1 if UMI *j* is derived from gene *i*. We also define the probability of generating a molecule from a particular gene according to the probability vector *ρ* (analogous to the nucleotide fraction in a typical bulk RNA-seq probabilistic model (Li and Dewey, 2011)). Hence, we can write the probability of observing a set of deduplicated UMIs 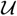 as follows:

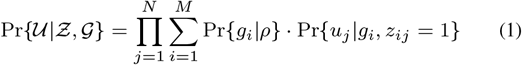

where 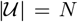 is the number of total molecules in the experiment (i.e. the number of deduplicated UMIs) and 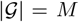 is the number of genes.

In this study, we take a variational Bayesian approach to gene-expression estimation. Therefore, instead of seeking the maximum-likelihood estimates, we infer (through variational approximation) the posterior distribution of *ρ*. This posterior distribution can be defined as:

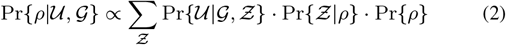

where both 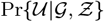 and 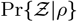 can be estimated via a variational approach (Hensman *et al.*, 2015). While numerous methods for expression estimation from bulk RNA-seq data have previously adopted a variational Bayesian approach (Nariai *et al.*, 2013, 2014; Hensman *et al.*, 2015; Patro *et al.*, 2017), they have all made use of uniform or uninformative priors. The novelty of our method comes from both adopting this approach in the single-cell context, and from setting the prior for *ρ*, in an informative, data-driven, and cell-specific manner. We expect that, subject to careful selection, information in a single-cell sequencing experiment can be meaningfully shared between distinct but related cells. Note that our method aims to accurately assign reads to the genes to which they map, and does not alter the expression level of genes with zero expression in the data, as may be the case with imputation-based approaches. We explain below how information from related cells, both within a sample and across assays, can be shared, and show how this principle can be applied under various scenarios to improve gene quantification accuracy.

### 2.2 Anchoring to obtain informative priors

Cells of the same type within a sample share similar expression patterns (Stuart *et al.*, 2019). However, due to both biological variability and, *crucially*, to the low capture rate and random sampling process in single-cell sequencing experiments, even cells of the same type do not always exhibit near-identical global gene expression profiles. This means that a given gene from two cells of the same cell-type within a sample could have varying expression estimates, and could be assigned different tiers in individual cells by the alevin algorithm. Specifically, a gene may be assigned tier 3 in one cell and tier 1 or 2 in the other, based on the *specific* sequenced reads and UMIs observed, and their mapping patterns. For example, if all of the reads arising from the gene come from an ambiguous region shared with other genes, then this gene will be assigned tier 3. Whereas if this gene, in another cell of the same type, has sequenced reads coming from a unique region, then it will be assigned as tier 1 (we have strong evidence of its existence in the cell). Hence, cells of the same type can potentially have different confidence levels in their gene estimates, irrespective of the associated count. This variation can be used to improve quantification of tier 3 genes. This scenario is depicted in Figure 2, which details how this information can play an instrumental role while quantifying these genes.

**Fig. 2:**
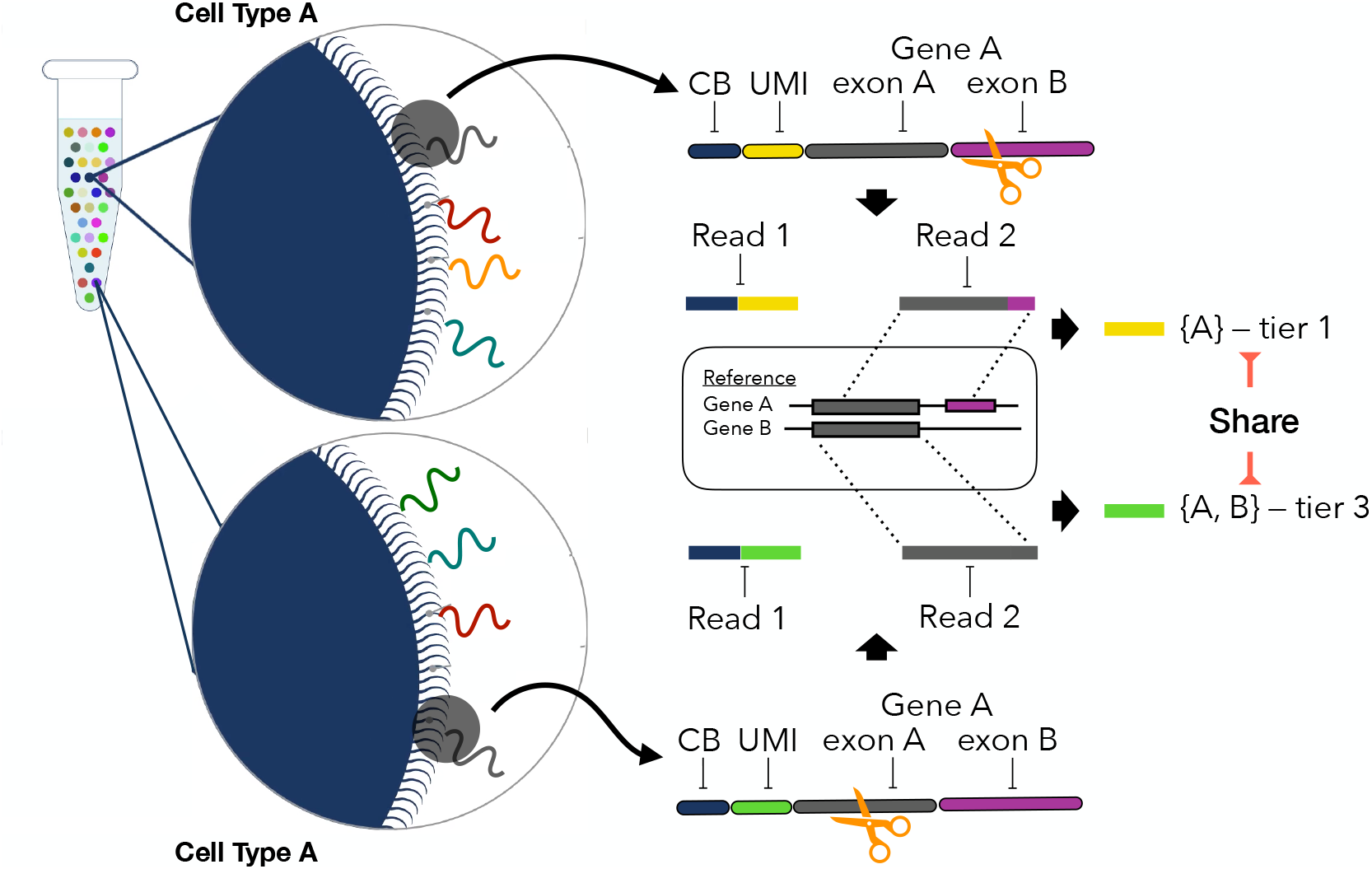
Motivation: Given two cells of similar cell type A, we select two reverse transcribed RNA molecules from Gene A, with unique Cell Barcodes (CBs) and Unique Molecular Identifiers (UMIs). Since the fragmentation of a molecule happens at random, the molecule from the first cell (on the top) is fragmented from a region uniquely identifiable for gene A while the molecule from the second cell (on the bottom) comes from sequence similar region of gene B. Top cell then has high confidence, tier 1 abundance estimate for gene A, while the bottom cell has a tier 3 estimate. Assuming the global expression profiles of these cells are similar, our proposed Bayesian model shares this information across cells to improve the quantification estimates for the second cell.

To first verify that it is possible to gain information in this way, we look at the fraction of cells that assign a particular gene to tier 3 out of the total number of cells where the gene is expressed. This is because priors will be informative only when obtained from cells where the gene has uniquely mapping reads (tier 1) or is influenced by reads mapping uniquely to genes sharing an equivalence class (tier 2). To do this analysis, we quantify the human PBMC 4k dataset (10x Genomics, 2017), using alevin supplemented with the whitelist output by Cell-Ranger. This experiment contains a total of 4340 whitelisted cells. The results of this analysis, shown in Figure 3, suggest that most genes are assigned tier 3 in less than 10% of the cells and, therefore, estimates from the other cells can be informative. For the 7484 genes that were assigned tier 3 in at least one cell, 37.1% are assigned tier 1 and 51.9% are tier 2 in other cells where the gene is expressed. Hence, the varying degree of confidence in expression estimates across cells can be leveraged in an informative way to improve tier 3 estimates. Note that all analyses henceforth are done using the 10x PBMC 4k dataset, except where mentioned.

**Fig. 3:**
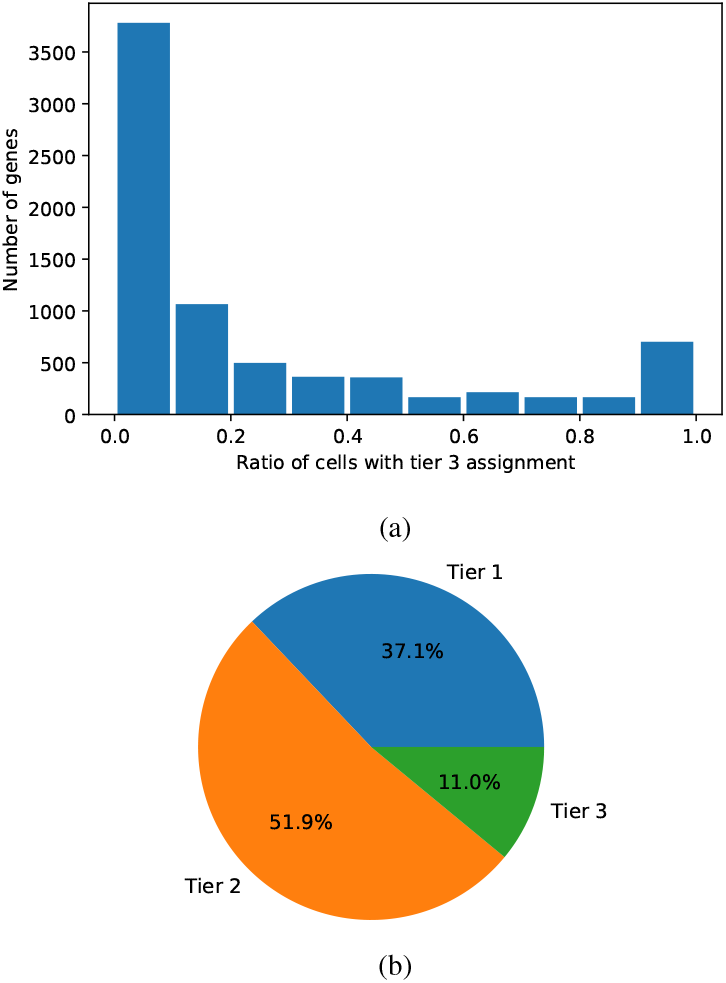
(a) The distribution of number of genes against the fraction of cells that have tier 3 assignment for these genes. For example, there are around 1000 genes that are assigned tier 3 in 0.1-0.2 fraction of the total number of cells. (b) The percentage of cells assigning each tier to the genes, showing that the degree of confidence in the quantification estimates varies across cells even for a single gene. Note that both these plots are made using 7484 genes that have been assigned tier 3 in at least one cell.

Based on these results, we can see that genes relegated to tier 3 in a given cell frequently have unique evidence in other cells within the same sample. To take advantage of this property, the next step is finding “neighboring” cells that might be useful for sharing this information to disambiguate read assignment between tier 3 genes. We only use information from cells with a similar global expression profile within the sample. To find similar cells for sharing this evidence, we use Seurat’s (Stuart *et al.*, 2019) cellular barcode anchoring scheme that defines a framework to connect two experiments based on the similarity in the gene expression patterns of the cells’ assayed in the two experiments. The algorithm works by calculating the *ℓ*2 distance across datasets, generating two distance matrices and then defines anchors as cells that are neighbors under both distance measures. The full algorithm implemented in Seurat is more involved, and includes various scoring metrics and parameters. Although initially intended for matching cells across samples, we use this anchoring algorithm to connect cells *within* the sample, in order to define cell-specific priors as input for our Bayesian algorithm.

To generate cell barcode anchors, we first quantify the sample using the standard alevin algorithm (henceforth referred to as EM), and divide the quantification estimates for all cells into two equal sets. We then run the Seurat anchoring algorithm on these sets, treating the two subsets as two separate samples. In order to identify anchors for a larger number of cells, we repeat the anchoring step multiple times, randomly dividing the quantifications into two equal groups each time. We repeat the anchoring step 30 times for all experiments in this manuscript, as we observe on the simulated data that the gain after 30 iterations is small (Figure 4). We filter the anchors based on the score output by Seurat, using only anchors with a score greater than 0.5. In a typical single-cell experiment, this is expected to find anchors for about 80% of the cells. The prior for a cell is then defined as the expression estimates, using the original EM based alevin run, of the cell assigned as the anchor. However, this process can eventually assign multiple anchors for a single cell. To compensate for this, we calculate the prior by taking the average of the expression estimates from all the anchors. This prior is used to optimize the quantification estimates of tier 3 genes with multi-mapping reads in the alevin pipeline, while keeping the prior uniform for tier 1 and 2 genes.

**Fig. 4:**
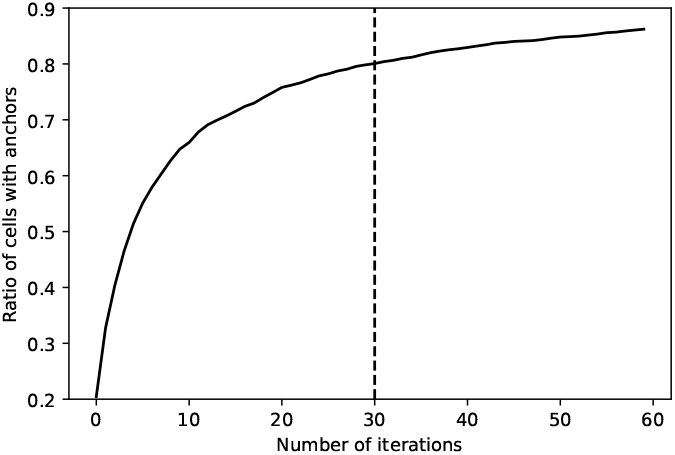
Ratio of cells matched in each iteration of the Seurat anchoring algorithm, splitting the dataset into 2 random, equal sets in each iteration.

## 3 Results

### 3.1 Improved estimates using intra-sample information

To test the hypothesis that combining the Bayesian framework with priors obtained from the anchoring procedure described in Section 2.2 can lead to improved quantification estimates for tier 3 genes, we devised two separate experiments. We detail these two setups below, one relying on simulated data, and the other relying on experimental data with “equivalence class knockout”.

#### 3.1.1 Simulated data

To analyze the improvements in gene quantification estimates on simulated data, we use the empirical dscRNA-seq data simulation tool Minnow (Sarkar *et al.*, 2019). Minnow models various features and protocols involved in the generation of dscRNA-seq data, like PCR amplification and sequencing errors, to generate fastq files with the reads and the true cell-by-gene count matrix. We use Minnow to simulate a dscRNA-seq experiment with 4340 cells and ~20 million UMIs using alevin (EM-based) quantifications on the 10x PBMC dataset as input. We then compared the quantification estimates against the truth, predicted on the simulated data using alevin, with and without priors, and Cell-Ranger. The priors from VBEM based alevin were generated, as explained above, using the Seurat anchoring algorithm iteratively.

The results from this analysis are presented in Figure 5(a), where VBEM represents quantification estimates using priors and EM signifies the quantification estimates without priors. We calculate the Spearman correlation between each method and the ground truth provided by Minnow, focusing on genes assigned tier 3 in individual cells by alevin. While the fraction of expressed genes assigned tier 3 in each cell is low, as shown in Figure 5(b), improvement in the accuracy of the gene abundance estimates is significant across hundreds of cells and shows that using informative priors, even from within a sample, can improve quantification. The result also shows that the correlation between estimates from Cell-Ranger and truth is much lower. This is expected since these genes will have a high number of multi-mapping reads that will be discarded, not just when using Cell-Ranger but also when using other dscRNA-seq quantification methods.

**Fig. 5:**
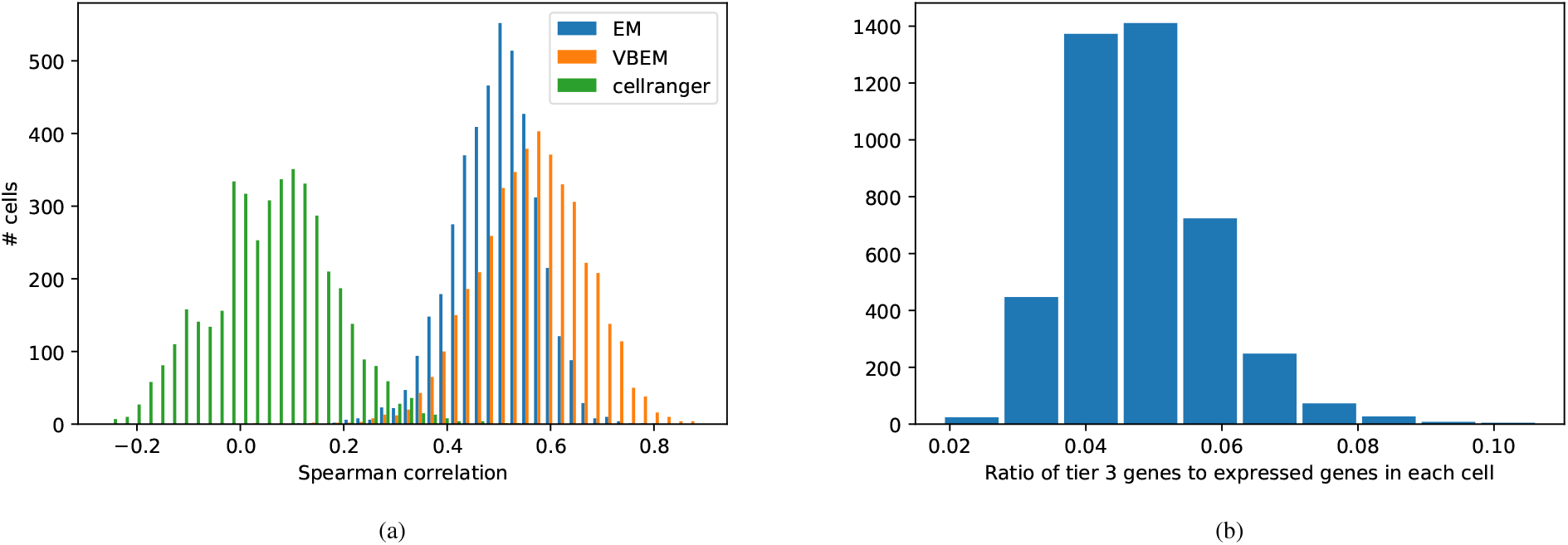
(a) Comparison of the cell-wise Spearman correlation of tier 3 genes quantified using Cell-Ranger, EM based alevin, and VBEM based alevin on simulated experiment. (b) Ratio of tier 3 genes in each cell (these are genes that *may* be impacted by the priors, leading to increased correlation with the truth).

#### 3.1.2 Experimental data with knockouts

To test that our proposed VBEM method, given informative priors, can improve the accuracy of experimental data quantification, we performed an experiment that we refer to as equivalence class knockout (KO). Alevin’s pipeline for dscRNA-seq quantification has multiple phases. After the initial phase of cell barcode whitelisting and read mapping, alevin outputs an intermediate file. This file contains details of the transcript equivalence classes, including the associated cell barcodes and UMI counts. These equivalence classes are similar to the gene-level equivalence classes explained before, except that the class labels are transcripts that share UMIs after the deduplication step. We observe, that a majority of the genes assigned to tier 1 and 2 after UMI deduplication are the ones associated with transcript equivalence classes of size 1 (labeled by a single transcript). In order to increase ambiguity in this data, we can remove all the transcript-unique equivalence classes from the intermediate file, with the expectation that this knockout will result in a number of genes migrating from tier 1 and 2 to tier 3, as demonstrated in Figure 6(a). In essence, by doing this, we are removing some of the read evidence that will eventually lead to high confidence gene abundance estimates in tiers 1 and 2. The impact of this on the distribution of tier 3 genes in the PBMC dataset is shown in Figure 6(b). This shows that the knockout process results in an increased number of genes that are assigned tier 3 across all cells. Note that the KO dataset will also have a smaller number of UMIs, because of the removal of unique equivalence classes from the intermediate file, but this will not impact our comparative analysis, as explained below. Also, note that we only knock out here equivalence classes that are *transcript-unique*, and that there will still be a considerable number of gene-unique equivalence classes after parsimonious UMI deduplication has taken place.

**Fig. 6:**
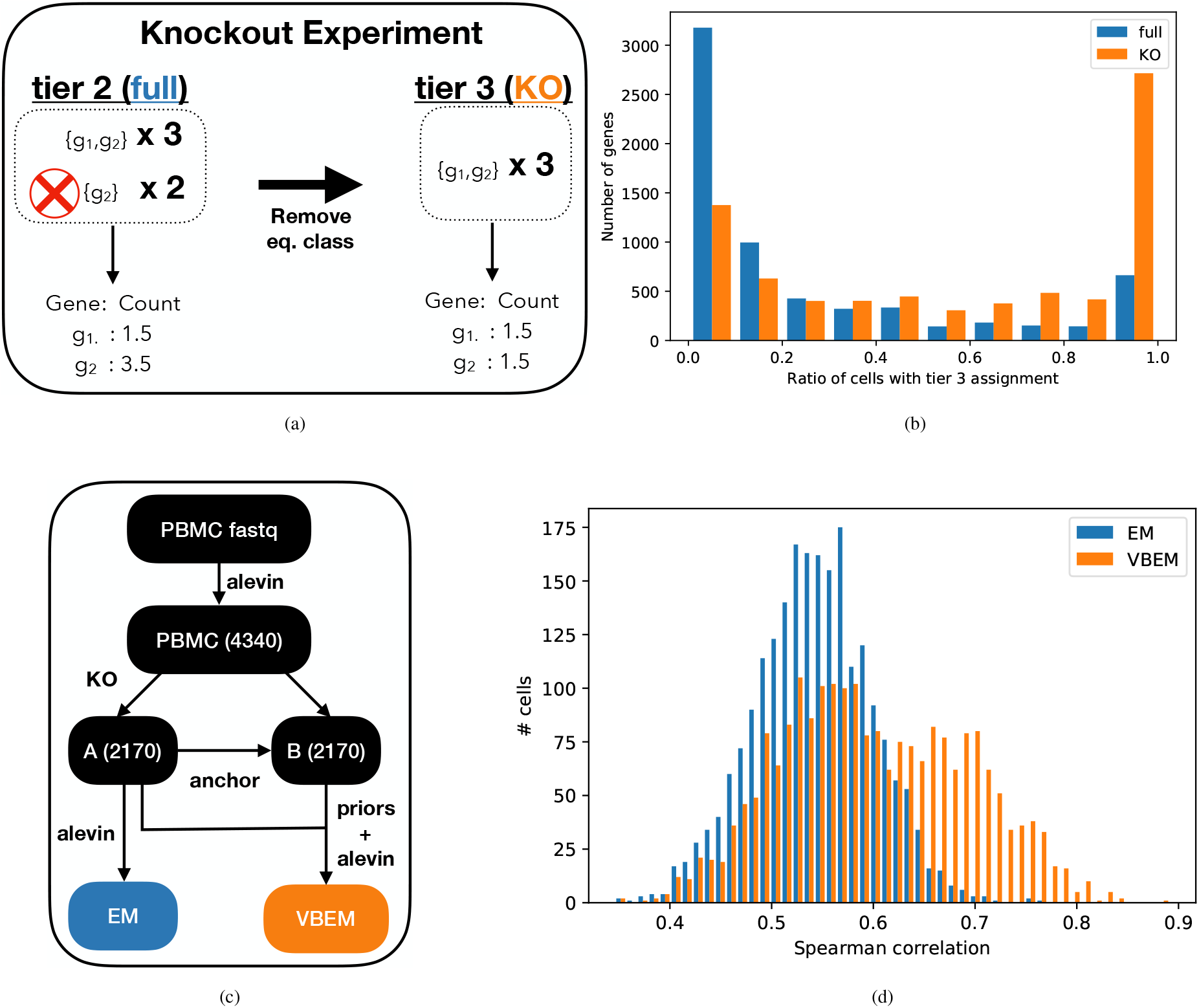
(a) A toy example explaining the knockout experiment. We use the equivalence classes from an alevin (EM based) run on the full PBMC dataset and remove all the transcript-unique equivalence classes to generate a knockout (KO) sample. In the example, assume each gene has a single transcript. (b) The distribution of number of genes against the fraction of cells that have tier 3 assignment for these genes both in the original dataset and the knockout dataset, with a shift towards more tier 3 assignments and increased ambiguity. (c) In the knockout framework, to validate the improved Bayesian approach, we designed the following pipeline. We quantify the full human PBMC dataset (4340 cells) cells and randomly divide the experiment in two equal parts (A and B). We knockout unique equivalence classes in set A (2170) cells and repeat the quantification step to generate EM based estimates. In parallel, we also quantify set A knockout dataset using Bayesian priors. These priors are learnt from set B, without the knockouts, and quantified initially using the EM approach. This gives us the VBEM estimates on set A for comparison. (d) Comparison of the cell-wise Spearman correlation for tier 3 genes from EM based alevin with VBEM based, prior enhanced alevin on real data with knockouts (removal of unique equivalence classes). This shows improved estimates under VBEM, with a higher correlation against the initial dataset, without knockouts.

In order to ensure that we have cells with high confidence quantification estimates to provide our KO cells with an informative prior, we did not perform the knockout procedure on the complete PBMC dataset. Instead, we took the alevin quantification estimates on the PBMC dataset, which has 4340 cells, and divided it into two sets containing equal numbers of cell barcodes. One of these sets, *A*, is our test set from which we knockout unique equivalence classes and the other set, *B*, is used to generate priors. Iteratively using the Seurat anchoring algorithm as before, we first find anchors for set *A* in set *B*, then obtain priors from set *B* and run the alevin VBEM quantification method. We also quantify the KO set *A* using the EM-based alevin method. These steps are outlined in Figure 6(c). Observe that there can be a bias in the tier assignment of genes that are anchors for tier 3 genes in the KO experiment. This is because we are removing equivalence classes only in set *A*. Hence, the ratio of anchor genes in set *B* that are assigned tier 1 and 2 in KO may be higher than in the real dataset. This can amplify the accuracy of the VBEM method in the knockout experiment, but will also reflect the actual gain possible for this methodology under varying circumstances, such as in samples with higher read depth. Note that we can not run Cell-Ranger on this dataset because it utilizes the intermediate file output by alevin, which can not be processed directly. However, we expect similar results as those observed in simulated data, since multi-mapping genes are not quantified by Cell-Ranger.

In our comparison between the two methods, we find the Spearman correlation for each cell between the original, EM based alevin estimates of the cells in set *A* and the estimates using the KO set *A* under each method. Because the original set *A* has more high confidence tier 1 and 2 genes, we expect the estimates to be of higher accuracy. The results from this analysis are presented in Figure 6(d), which shows that the cell-wise correlations of the VBEM predicted abundances on the KO dataset are higher compared to the original estimates than are the EM estimates on the KO dataset. Note that these correlations are calculated for genes that are assigned tier 3 in the KO set *A*, since those are *the only genes impacted by the priors*. This test shows that utilizing the anchoring procedure and extracting informative priors, combined with using a VBEM based quantification procedure, can lead to higher accuracy in abundance estimation.

It is also interesting to note that the anchoring scheme finds high scoring anchors between set *A* and set *B* for only 934 cells. The effect of this limited anchoring shows up in the correlation histogram as a bimodal distribution in the VBEM correlation values, signifying that, as expected, only some of the cells — those for which we were able to find an anchor in the set *B* — have improved correlation with the original quantification estimates.

#### 3.1.3 Information sharing does not affect rare cell types

A common concern when sharing information across cells in single-cell RNA sequencing analysis is that it may contribute to loss of heterogeneity among the quantified cells (Huang *et al.*, 2018; Andrews and Hemberg, 2018), removing not only technical “noise”, but also important biological variability that leads to the detection of important features, such as rare cell types. To test the hypothesis that the proposed Bayesian framework does not “over-regularize” and lose rare cell types in downstream processing, we perform the following experiment. We use the human PBMC dataset with 10k cells (10x Genomics, 2018) and quantify the cells with both the EM and VBEM-based approaches, where, for the VBEM-based approach we use the same procedure of generating priors as discussed in Section 3.1.1. Next, we perform Seurat (Satija *et al.*, 2015) based clustering on the estimates generated from both the approaches separately and compare the clusters.

In Figure 7, we show the 2D UMAP embeddings of the clustered data, colored by cell-type annotations generated using marker genes, as detailed in the Seurat pipeline (Stuart *et al.*, 2019). We observe that the clusters with relatively smaller number of cells, such as pDC, Megakaryocytes and Dendritic cell, are not lost by the Bayesian correction method. In Table 1, we show that the number of cells is almost always preserved in the most abundant cluster of each cell type across the two quantification approaches. We also observe that CD14+ Monocytes and CD8 effector cell types are divided into two subclusters when quantified with EM while they are correctly identified as one in case of VBEM.

**Table 1.** The number of cells observed across various cell types is similar when clustering is performed on EM and VBEM based quantification estimates, suggesting that the information sharing approach does not eliminate meaningful heterogeneity in gene expression between cells. The annotations are generated using Seurat’s marker gene analysis.

| Cell Type / # cells | EM | VBEM |
| --- | --- | --- |
| CD16+ Monocytes | 318 | 322 |
| CD8 effector | 222 + 144 | 358 |
| CD4 Naive | 1015 | 1021 |
| Megakaryocytes | 49 | 49 |
| NK cell | 517 | 522 |
| CD14+ Monocytes | 1758 + 1211 | 2962 |
| pDC | 68 | 68 |
| CD8 Naive | 333 | 331 |
| B cell progenitor | 455 | 453 |
| Dendritic cell | 74 | 74 |
| CD4 Memory | 1428 | 1416 |
| Double negative T cell | 587 | 583 |
| pre-B cell | 916 | 925 |

**Fig. 7:**
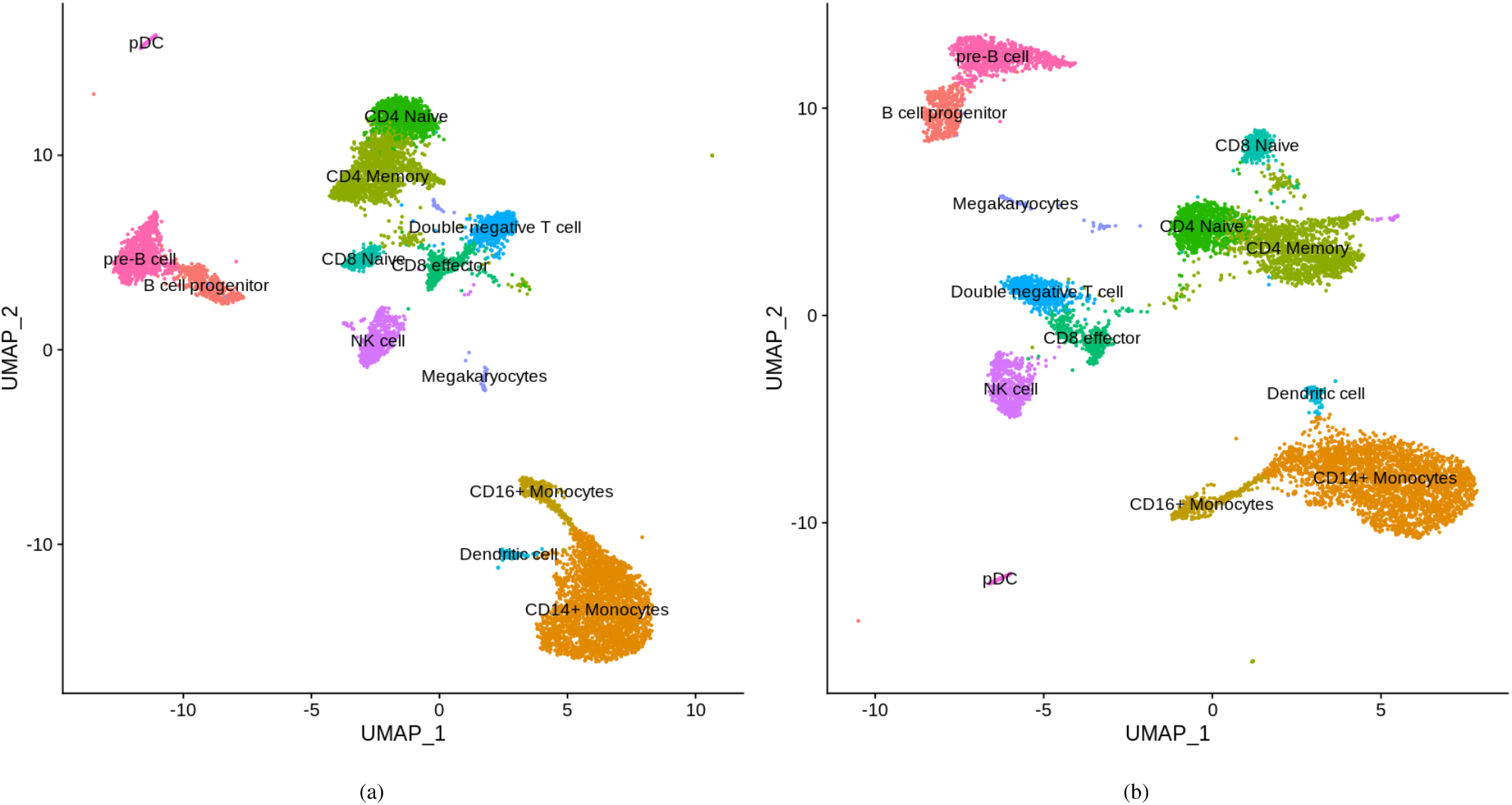
We perform cell clustering using Seurat on PBMC 10k dataset quantified using the EM (left) and VBEM (right) approaches in alevin and color the cells based on their cell type annotation generated using marker genes.

### 3.2 Improved estimates using multi-modal information

#### 3.2.1 Spatial Transcriptomics Data

Advancements in spatial transcriptomics (ST) have enabled scientists to relate cells with their location within a tissue. Specifically, it has been shown how combining ST with gene expression profiling in cancer data helps understand multiple components of tumor progression and therapy outcomes (Thrane *et al.*, 2018). 10x Genomics Visium is another interesting assay that provides higher resolution and throughput for spatial gene expression analysis. We use the open dataset provided by 10x Genomics of the fresh frozen mouse brain tissue with 2698 spots in the tissue and process the raw reads through the alevin framework to generate a gene count matrix for each spot.

To test the Bayesian framework of alevin, we simulate 2698 cells using the gene count matrix generated by processing the mouse brain ST visium data from 10x Genomics (2019). We first run EM based alevin on the simulated data and use the spatial 2D coordinates from the ST data to learn the prior, i.e. for each cell we use the nearest 8 cells and their mean gene expression from the EM estimates to generate the prior matrix. Then, we provide alevin with the prior matrix to re-quantify the simulated data using the Bayesian method to generate VBEM based estimates. In Figure 8 we show the cell-wise Spearman correlation of tier 3 gene estimates for both EM and VBEM based methods. We observe a global shift in the VBEM quantified data, reflecting the increased accuracy obtained using informative priors from cells located spatially close together. This result is particularly interesting, as it suggests that the empirical Bayesian framework we have introduced is modular and flexible, in that the generation of an informative prior is not tied to a specific procedure (e.g. the Seurat-based anchoring). Rather, the prior can be informed by data in the same sample, by assay-specific information (nearby or differential cell clusters in spatial data as also shown by Äijö *et al.* (2019)) and, perhaps, even across distinct modalities (e.g. between ATAC-seq and RNA-seq for cells assayed with both protocols in the same sample).

**Fig. 8:**
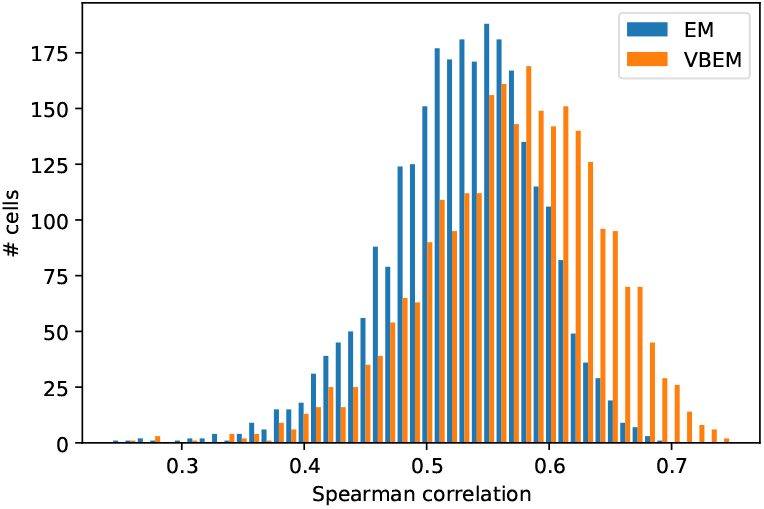
Comparison of the cell-wise Spearman correlation of tier 3 genes quantified using EM based alevin and VBEM based alevin on spatial transcriptomics data.

## 4 Discussion and Conclusion

In this work, we improve upon our previously proposed alevin pipeline for quantification of dscRNA-seq data. The existing alevin pipeline uses a maximum likelihood based procedure after the UMI deduplication phase to accurately resolve multi-mapping reads, which are typically discarded by other methods. While this approach uses unique read evidence from within a cell to optimize read assignment, it uniformly divides read counts where no unique evidence is available. The set of genes with this uniformly divided distribution are assigned tier 3 by in the alevin output. Our proposed method uses a Bayesian framework to improve tier 3 gene quantification.

This method works by sharing high confidence quantification information between cells. Information is shared only across cells that have similar gene expression profiles (or which are spatially proximate in the case of spatial transcriptomics data), but the exact expression estimates vary due to sparsity and uneven RNA capture in single-cell sequencing protocols. We show that, under several different experimental setups, our information-sharing framework consistently improves tier 3 dscRNA-seq quantification estimates. This approach is especially useful for highly ambiguous estimates where there is no intra-cellular unique information available to accurately quantify the genes, but where simply discarding the multi-mapping reads would lead to the loss of potentially important biological information.

While we have focused on tier 3 genes in this study, this information sharing model can be extended further to improve the UMI deduplication procedure as well, before the construction of equivalence classes. For example, instead of basing UMI deduplication on the principle of parsimony in alevin, priors can be used to drive deduplication. This can lead to improvements in abundance estimates for all genes in the reference. Similarly, with advances in single-cell sequencing protocols, this framework can be extended to incorporate priors from different technologies. For example, as we have demonstrated, spatial data can be useful for setting the prior in the proposed alevin framework. This improves accuracy by relying not on similar gene expression profiles, but cells that are in close proximity in physical space. Further, one can imagine that other assays, like paired single-cell ATAC-Seq and RNA-seq, would allow useful information sharing within the same sample but across data types and modalities. We believe this framework has the potential to open a new direction of enabling multi-modal information sharing to improve quantification of single-cell data.

## Author’s contributions

AS, LM, HS, and RP devised the methods. AS, LM, and RP implemented the methods. AS, LM, and RP devised and carried out the experiments. All authors wrote and approved the manuscript.

## Acknowledgments

The authors would like to thank Michael Love, Rahul Satija, Tim Stuart and Charlotte Soneson for fruitful discussions surrounding some of the approaches taken in this manuscript.

## Resources

The pipeline to replicate the analysis can be found at https://github.com/COMBINE-lab/alevin-paper-pipeline/tree/master/bayesian_alevin. We used the gencode 28 reference for human and gencode mm10 for the mouse references. We use Seurat version 3.0.2 and cellranger version 3.1 with the following commands:

1. index: cellranger mkref ––genome=ref ––fasta=genome.fa ––genes=genes.gtf ––nthreads=16
2. quantification: cellranger count ––id=cellranger ––fastqs=fastqs ––localcores=20 ––localmem=120 ––transcriptome=ref

